# The canine respiratory epithelium is a permissive ecosystem for influenza interspecies transmission and emergence

**DOI:** 10.64898/2026.06.04.730051

**Authors:** Hanting Chen, Jack Hassard, Jiayun Yang, Callum Magill, Toby Carter, Jean-Remy Sadeyen, Aimi Ito, Clio Duerr, Hannah Montgomery, Savitha Raveendran, Kieran Dee, Maximilian N.J. Woodall, Grace B. Tyson, Maria M. Afonso, Verena Schultz, Claire M. Smith, Margaret Hosie, Stuart M. Haslam, Munir Iqbal, Pablo R. Murcia

## Abstract

The outcome of virus spillover ranges from dead-end infections to pandemics and is underpinned by host–pathogen interactions as well as evolutionary and epidemiological processes. The emergence of novel influenza A viruses (IAVs) has been associated with reassortment events involving multiple species, highlighting the importance of reservoir and intermediate hosts in viral emergence. Highly pathogenic H5N1 IAVs of the 2.3.4.4b genotype have caused a panzootic affecting a broad range of mammals. The role of dogs—arguably the most popular companion animal and a natural host of IAVs—in the ecology of IAVs under this new zooepidemiological scenario is unknown. To address this, we characterised the glycome of the dog respiratory epithelium, infected canine tracheal explants with multiple IAVs (including canine H3N2 and H3N8, equine H3N8, avian H3N8 and H5N1, swine H1N1, human H1N1 and H3N2, and bovine H5N1 viruses), and determined their cellular tropism. We show that the respiratory tract of dogs presents abundant sialylated glycans known to act as IAV receptors. Further, most IAVs (including 2.3.4.4b viruses) infected and replicated in dog tracheas, targeting mainly ciliated cells. Serological testing showed evidence of influenza spillover infections in dogs from the UK. Overall, our results show that the canine respiratory tract can provide a suitable environment for the generation of new IAVs. Given the multi-host contact networks of dogs in nature, they could act as recipients, bridging hosts, and/or mixing vessels for multiple IAV lineages, playing a central role in the ecology of influenza emergence.

## INTRODUCTION

Virus spillover (i.e. interspecies infections) poses a constant threat to humans and animals. From a public health perspective, they can lead to outbreaks, epidemics and pandemics, while from a veterinary standpoint they can affect animal health and trade, as well as food security. Further, the spread of emerging viruses among wildlife can result in biodiversity losses.

Although humans and animals are constantly exposed to a myriad of viruses circulating among sympatric species, viral emergence is relatively uncommon. Indeed, the outcome of spillover infections is highly variable: while SARS-CoV-2, influenza A viruses (IAVs) and human immunodeficiency virus (HIV) caused pandemics and established as human endemic viruses^1–3^, Nipah virus and Middle East respiratory syndrome coronavirus caused limited outbreaks^4,5^. In turn, Hendra and rabies virus cause single dead-end human infections^6,7^. For respiratory viruses such as IAVs, the probability of infection during spillover exposure will be affected by multiple host barriers that are specific to individual virus/host combinations^8^. These include, among others, the presence and composition of mucus, the availability of functional receptors on the cell surface, the pH of the endosome, and the effectiveness of the antiviral innate immune response.

Ecological and evolutionary factors play a central role in the outcome of spillover infections. As IAVs possess a segmented genome and exhibit a relatively broad host range, spillover can result in viral coinfections that lead to segment reassortment, an evolutionary process that can increase the fitness of otherwise poorly adapted viruses. Indeed, virus reassortment has been associated with the generation of pandemic IAVs^3^.

The ecology and epidemiology of H5N1 IAVs have changed dramatically in recent years. This lineage has spread globally in wild birds, spilling over into numerous mammalian species including mink, racoon dogs and sea lions^9^. One of the most striking phenomena of the H5N1 panzootic is the emergence of the 2.3.4.4b genotype in dairy cattle^10^. While previous studies showed serological evidence of IAV infection in cattle^11^ and cattle are susceptible to infection under experimental conditions^11^, the absence of bovine IAV lineages led to the notion that cattle could not become reservoirs of IAVs.

Similarly, dogs had also been perceived as a species refractory to influenza emergence. This perception changed in the twenty first century after the emergence of three novel canine influenza virus (CIV) lineages: an equine-origin H3N8^12^, an avian-origin H3N2^13^, and a swine-origin H1N1^14^. Currently, H3N8 CIV is considered extinct^15^, and only H3N2 and H1N1 CIVs are circulating.

Dogs are one of the most popular domestic animals^16^. They are bred as pets, used for sports (racing and hunting), work (e.g. policing and herding) and farmed for meat. Consequently, they interact closely with other domestic animals, wildlife, and humans. Due to this these frequent cross-species interactions, dogs can act as a source of zoonotic viruses (e.g. rabies) and as recipients of anthropozoonotic viruses such as SARS-CoV-2^17^, IAV^18^ or monkeypox virus^19^. Notably, spillover of canine distemper virus (CDV) from dogs was shown to be the main driver of CDV infections in lions in Africa^20^, highlighting the significance of the interface between domestic and wild animals. Dog population structure varies across urban and rural environments and is shaped by regional ownership practices, husbandry systems, and cultural norms.^16^. Studies on rabies and canine influenza have shown that the structure and contact network of the host population is central to virus transmission^21,22^. In addition, exposure to viruses of humans or animals will depend on the ecological context. For example, pet dogs in many countries will spend most of their time indoors in close contact with humans, while farm dogs will be regularly exposed to other domestic animals. In contrast, dogs used in hunting sports or living in rural free-roaming populations will have high contact rates with wildlife.

With regards to influenza, dogs are susceptible to infection by IAVs derived from humans and animals. Serological studies have shown evidence of dog exposure to human H1N1 and H3N2 viruses^23,24^, and experimental studies using live animals or canine-derived tissues have shown that the canine respiratory tract is susceptible to infection by IAVs of different origin^25–27^.

Given the dynamic nature of the current H5N1 panzootic and the complex ecology and evolution of IAVs, we aimed to determine whether the dog respiratory tract would be a suitable environment for the generation of IAVs with potential for zoonosis and interspecies transmission. We used canine tracheal explants to characterise the glycomic profile of the dog trachea and determine the cellular tropism and growth kinetics of IAVs of different origins that included evolutionarily distinct human viruses, canine, equine, swine, and avian IAVs, as well as avian and bovine H5N1 2.3.4.4b viruses. We also assessed IAV exposure levels by testing serum samples collected between 2022 and 2024 in the UK.

## RESULTS

### Canine tracheas contain glycan receptors terminating in both α2-6 and α2-3 linked sialic acid

To determine the types of sialylated glycans (*i.e* potential IAV receptors) in the canine respiratory tract, we conducted detailed glycomic analysis of canine tracheal tissue and compared the results to cultured trachea explants, which were used as a model system for IAV infection. N-Glycans terminating in sialic acid, predominantly N-acetylneuraminic acid (NeuAc), were abundant in the canine trachea (Figure 1A), particularly glycans at *m/z* 2792, 2809, 2966, 3603 and 3777. These sialylated glycans ranged from mono- and di-sialylated bi-antennary glycans (e.g. *m/z* 2605, 2792 and 2966) to tetra-sialylated, tetra-antennary glycans (e.g. *m/z* 4587, 4761, 5006 and 5211). We detected structures containing sialyl-lewis X/A structures (NeuAcα2-3Galβ1-3/4(Fucα1-3/4)GlcNAc), consisting of sialylated N-acetyllactosamine (LacNAc, NeuAc-Gal-GlcNAc) branches with an additional fucose residue (e.g. *m/z* 4196, 4645, 4761, 4819, 5006, 5064, 5211). In addition to sialic acid, glycans terminating in Galα1-3Gal (e.g. *m/z* 2652 and 2809) and ABO blood group A epitopes (m/z 3024, 3964, 4819, 5064) were also detected. The cultured trachea explants were highly representative of the canine trachea tissue and expressed similar glycan structures with minor differences in their relative abundances (Figure 1B). To establish the ratio of “human” vs “avian” influenza virus receptors we treated the glycans with sialidase-S, which specifically cleaves α2-3 linked sialic acid (Supplementary Figure 1). This revealed that both the canine tracheas and the cultured trachea explants contained an approximately equal mix of both α2-6 and α2-3 linked sialic acid (Figure 1C), which could be preferentially bound by human and avian influenza viruses, respectively.

**Figure 1.**
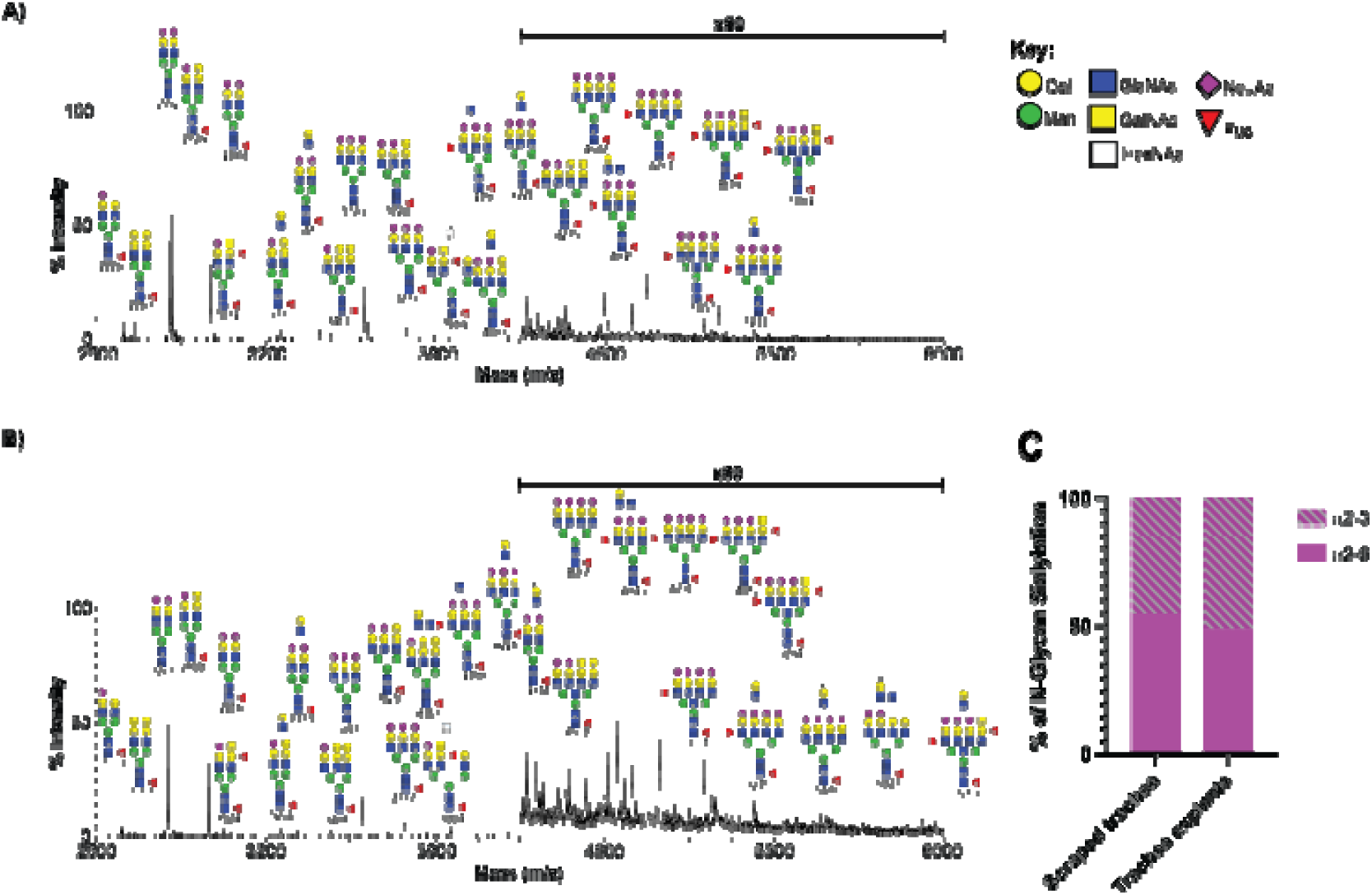
Canine tracheas contain a mix of ⍰2-3 and ⍰2-6 linked sialic acid receptors. Annotated MALDI-TOF spectra of permethylated N-glycans from **A)** canine trachea and **B)** cultured canine trachea explants. Annotations show [M + Na]^+^ molecular ions. Peak annotation is based on composition, biosynthetic knowledge and MS/MS analysis. **C)** Proportions of ⍰2-3 and ⍰2-6 linked sialic acid in canine trachea and cultured trachea explant N-glycans.

The majority of O-glycans from both the trachea and trachea explants contained ABO blood group A epitopes, however molecular ions corresponding to sialylated core 1 and core 2 structures were detected in both (Supplementary Figure 2). Sialic residues were primarily linked to the initial N-acetylgalactosamine (NeuAcα2-6GalNAc) and/or to the galactose (Gal) on the β1-3 branch (NeuAcα2-3Galβ1-3GalNAc). The relative abundances of sialylated O-glycans were higher in the trachea explants and additional molecular ions (m/z 1345 and 1706) were detected, corresponding to mono- and di- sialylated core 2 O-glycans. For these structures, the sialic acid on the β1-6 branch could be either α2-3 or α2-6 linked (NeuAcα2-3/6Galβ1-6GalNAc).

To examine the distribution of sialic acids *in situ* we stained formalin-fixed, paraffin-embedded canine tracheal explants with lectins that bind specifically to α2-3- or α2-6-linked sialyl glycans (*Maackia amurensis* [MAL] and *Sambucus nigra* [SNA], respectively). MAL staining was predominantly observed on the apical surface of the epithelium, and SNA staining was observed in the submucosal glands as well as on the apical and intraepithelial regions of the trachea (Figure 2). Overall, our results show that receptors for avian and human IAVs are broadly distributed across accessible regions of the canine respiratory epithelium.

**Figure 2.**
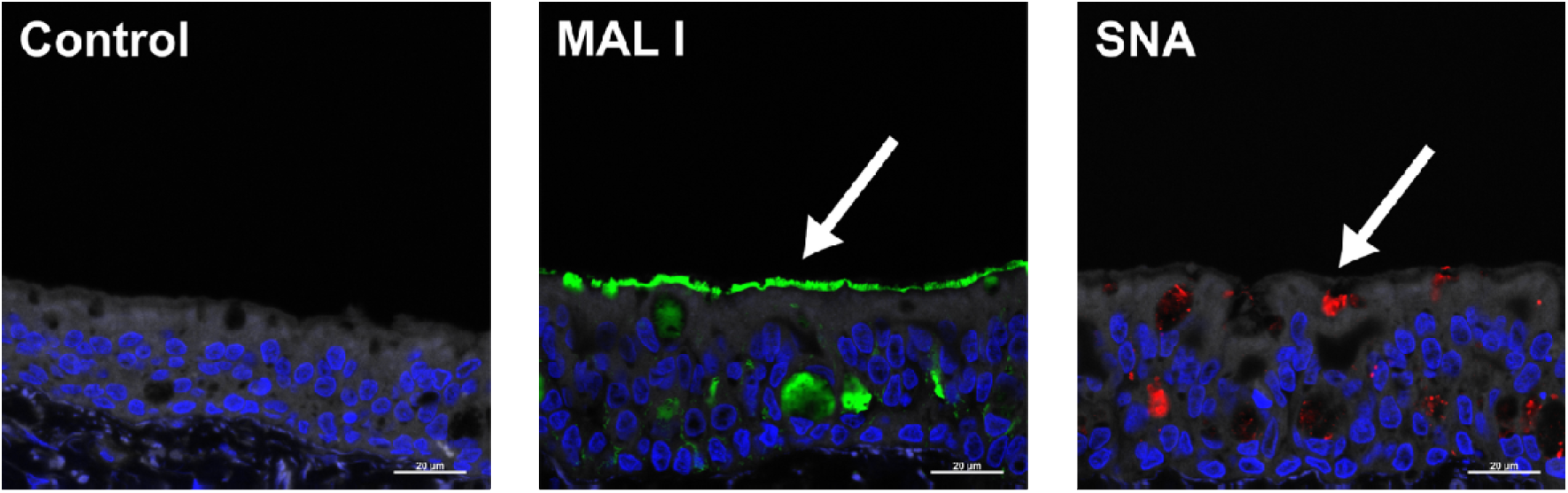
Staining of canine tracheal explants with lectins. MAL-I (green) and SNA (red) staining of formalin-fixed, paraffin-embedded canine tracheas. Scale bar: 20 μm. Arrows indicate cells displaying positive staining.

### Host-specific IAV lineages exhibit distinct replication kinetics in dog tracheas

We infected canine tracheal explants with a broad range of IAVs derived from different hosts to compare their replication fitness in the canine respiratory epithelium. Initially, we infected explants with biosafety-level 2 viruses that included H3N2 and H3N8 CIVs (CIV[H3N2] and CIV[H3N8] respectively); a low-pathogenic H3N8 avian influenza virus (AIV[H3N8]); H3N8 equine influenza virus (EIV[H3N8]); and two human isolates, H1N1 (IAV[H1N1]) and H3N2 (IAV[H3N2]). To assess the replication kinetics of H5N1 bovine influenza virus (BIV[H5N1]) and a highly pathogenic, genotype 2.3.4.4b avian influenza virus (AIV[H5N1]) in dog tracheas, tissue explants were prepared and infected with the viruses in high containment laboratories in separate experiments using the same protocols. We used a low dose of 200 plaque forming units (PFUs) that is likely to mimic natural infection conditions and capture differences in infection phenotypes that could be masked by higher, saturating doses. Viruses were titrated every 24 hours until 5 days post-infection. IAV(H3N2) was the only virus that did not replicate in dog tracheas. All the other viruses replicated consistently. CIV(H3N2) and IAV(H1N1) displayed the highest levels of replication, with peaks of ∼10^7^ PFU/ml (Figure 3). Both 2.3.4.4b viruses, BIV(H5N1) and AIV48(H5N1), also replicated in dog tracheas, reaching peak titres of ∼10^4^ PFU/ml (Figure 4). Overall, our results indicate that the canine respiratory epithelium is susceptible to infection by a broad range of IAVs derived from different host species.

**Figure 3.**
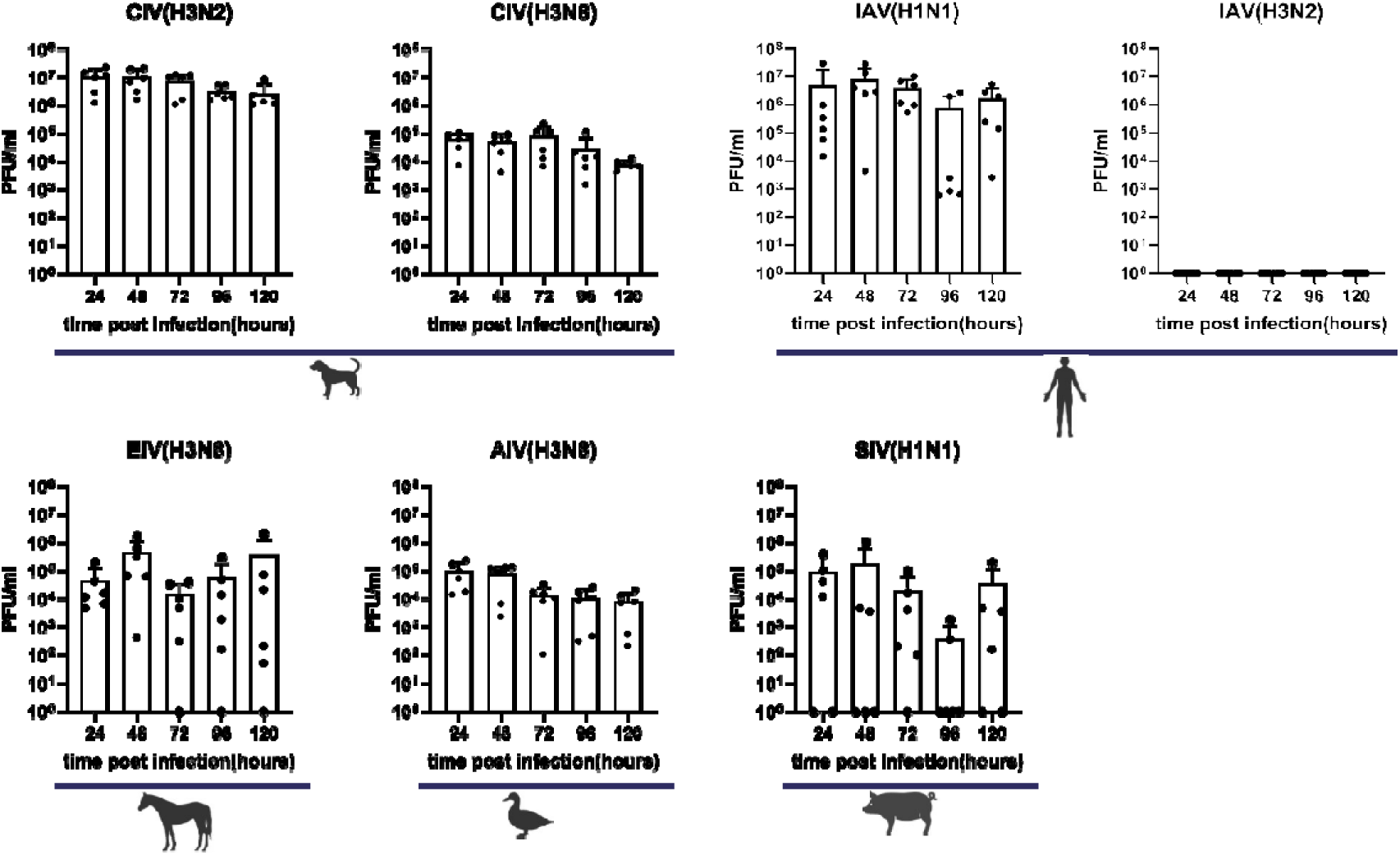
Growth kinetics of IAVs of different origin in dog tracheal explants. Replication kinetics of CIV(H3N2), CIV(H3N8), AIV(H3N8), IAV(H1N1/2018), IAV(H3N2/2018), EIV(H3N8), and SIV(H1N1) in explants derived from canine tracheas. Viral titres at individual timepoints are shown in white columns as solid black dots. Each dot represents an individual measurement, and bars represent the mean of six values. Data are combined titres from three independent experiments.

**Figure 4.**
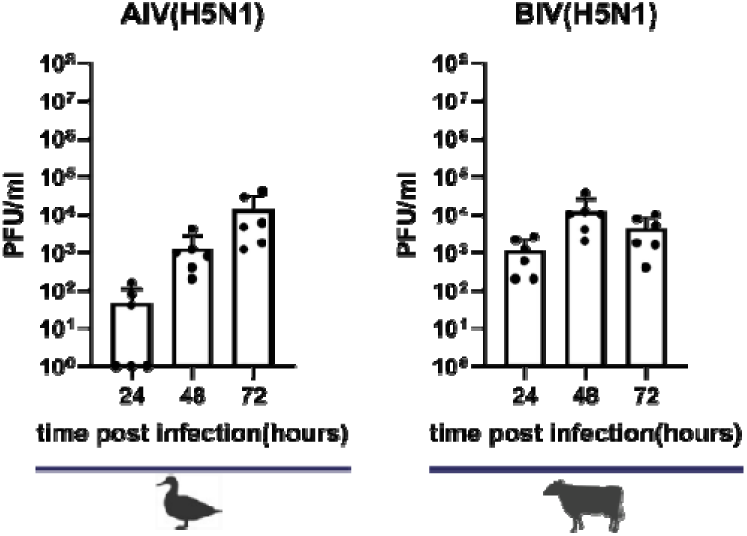
Growth kinetics of H5N1 bovine influenza virus (BIV[H5N1]) and a highly pathogenic, genotype 2.3.4.4b avian influenza virus (AIV[H5N1]) in explants derived from canine tracheas. Viral titres at individual timepoints are shown in white columns as solid black dots. Each dot represents an individual measurement, and bars represent the mean of six values. Data are combined titres from three independent experiments.

### IAVs derived from different hosts share similar cellular tropism

To define the cellular tropism of different IAV strains, we performed double immunofluorescence staining using an antibody against viral nucleoprotein (that detects infected cells) together with antibodies against markers of ciliated (beta tubulin), goblet (MUC5AC), and basal (p63) cells. All IAVs -with the only exception of IAV(H3N2)- infected ciliated cells (Figure 5 and Supplementary Table 1). Detection of infected goblet cells was less frequent and displayed marked variation between donors (Figure 6 and Supplementary Table 2). However, it should be noted that goblet cells constitute a minority in the tracheal epithelium (Supplementary Table 2). Infection of goblet cells by BIV(H5N1) was not detected. Basal cells were rarely infected (Supplementary Figure 3). These results suggest that ciliated cells (the predominant cell type within the airways) and goblet cells could support coinfection with different IAVs, potentially providing a suitable cellular environment for reassortment.

**Figure 5.**
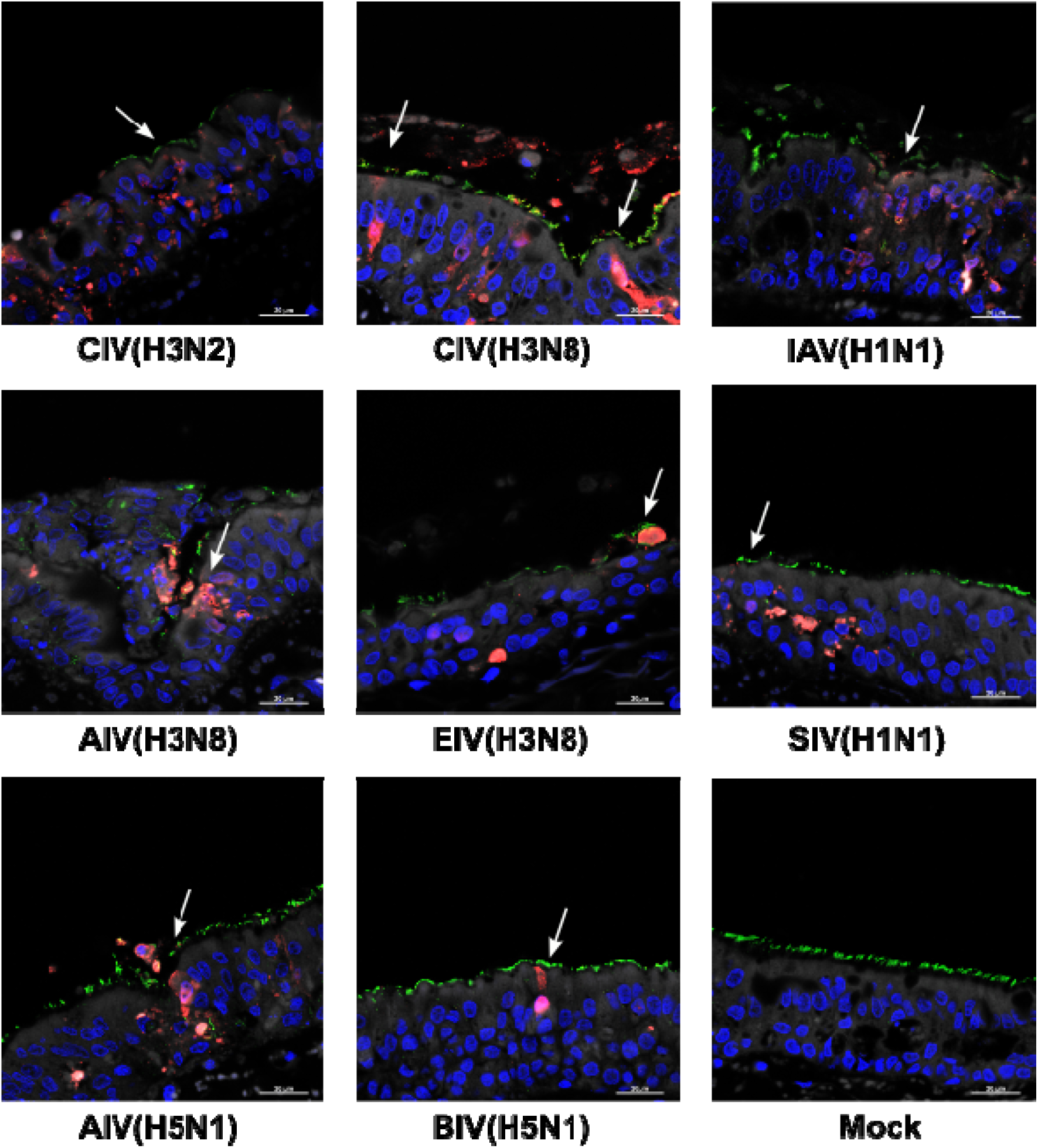
Cellular tropism of IAVs. Representative images of immunostained sections derived from canine tracheal explants infected with CIV(H3N2), CIV(H3N8), IAV(H1N1), AIV(H3N8), EIV(H3N8), and SIV(H1N1), AIV(H5N1), BIV(H5N1) and mock-infected. NP is shown in fluorescent red and indicates infected cells, whereas fluorescent green shows immunostaining of beta tubulin, indicating ciliated cells. Nuclei are stained with Hoechst and shown in fluorescent blue. Arrows indicate infected ciliated cells.

**Figure 6.**
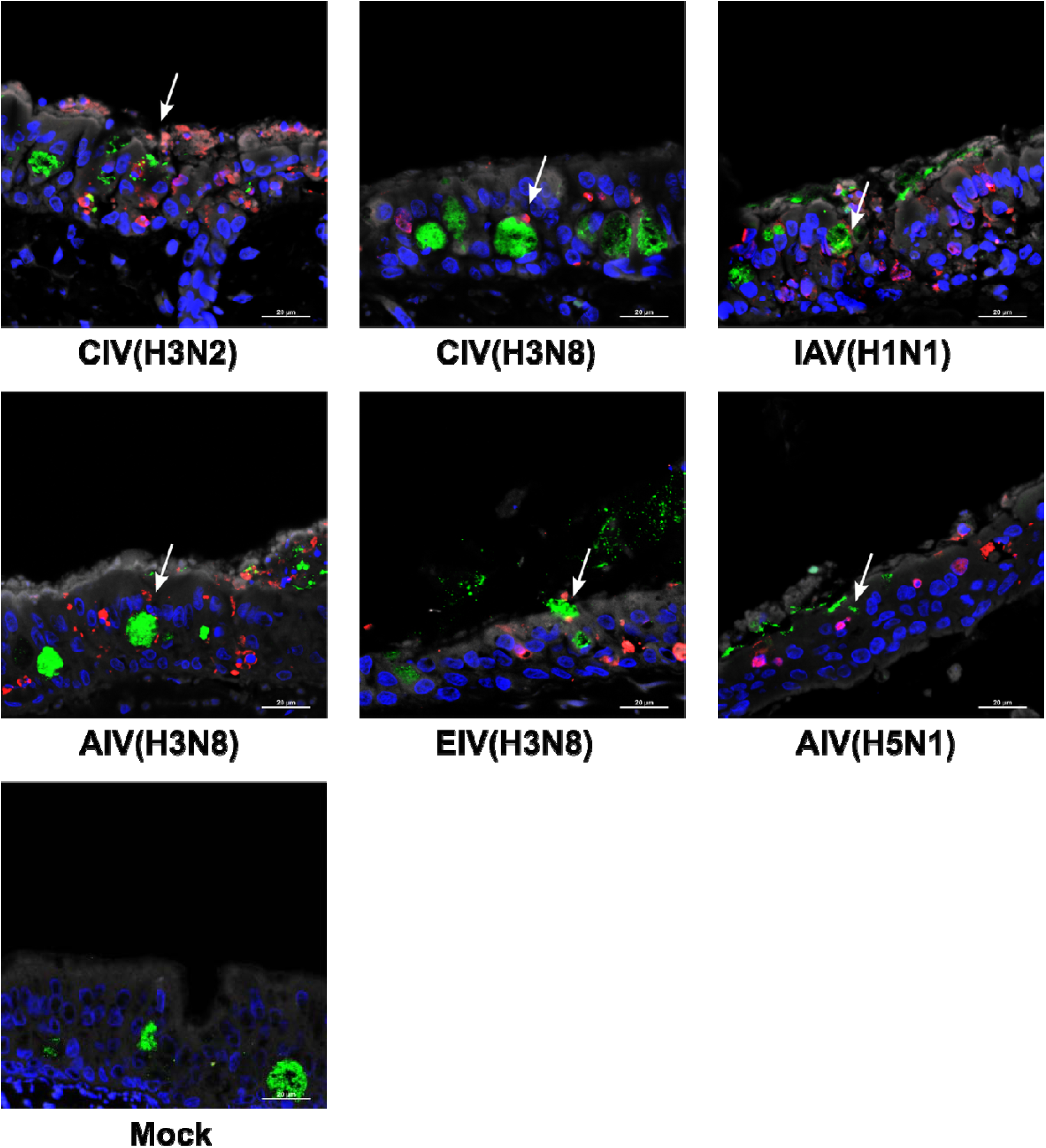
Cellular tropism of IAVs. Representative images of immunostained sections derived from canine tracheal explants infected with CIV(H3N2), CIV(H3N8), IAV(H1N1), AIV(H3N8), EIV(H3N8), AIV(H5N1), and mock-infected. NP is shown in fluorescent red and indicates infected cells, whereas fluorescent green shows immunostaining of MUC5, indicating goblet cells. Nuclei are stained with Hoechst and shown in fluorescent blue. Arrows indicate infected goblet cells.

### Influenza virus infection triggers widespread innate immune responses across the respiratory epithelium

For reassortment to occur two viruses must infect the same cell. However, secondary viral infections can be hindered by IFN-mediated responses elicited by the virus that causes a primary infection. We first assessed whether the IAVs under investigation triggered an innate immune response in the canine respiratory epithelium. Figure 7A shows immunostaining of infected (and control) explants with antibodies directed against Mx1, ISG15, and IFITM3, which are highly upregulated upon IFN stimulation and are broadly used as markers of innate immune activation^28,29^. Mx1 was highly expressed across the entire epithelium in explants infected with all viruses, whereas expression of the other two IFN-stimulated genes (ISGs) was variable (Figure 7B). Interestingly, Mx1 was highly expressed in explants infected with BIV(H5N1). Furthermore, IFITM3 expression was higher under basal conditions than in infected explants (Figure 7B). These results suggest that IAV infection triggers strain-specific epithelial responses.

**Figure 7.**
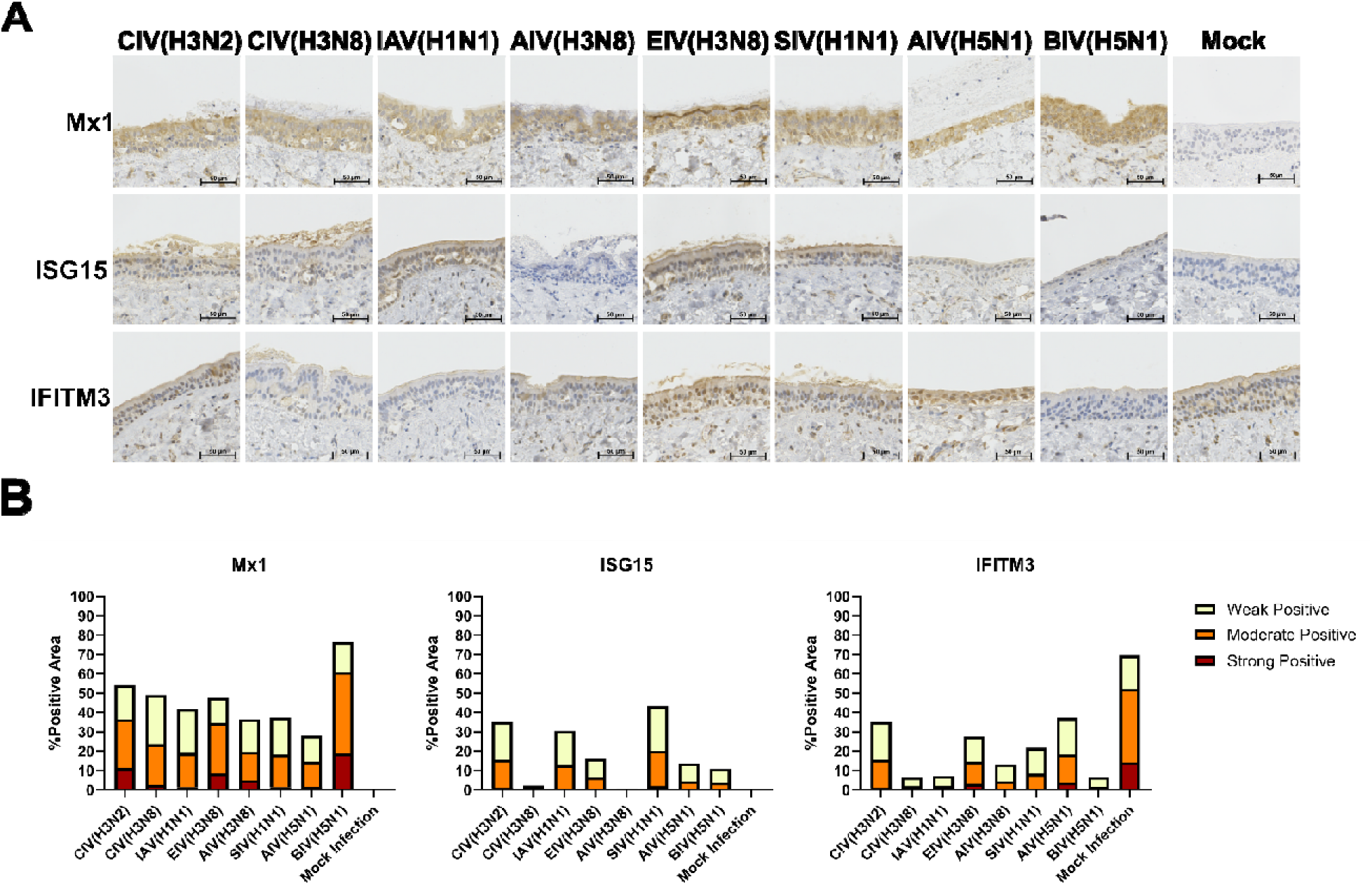
A. Expression of Mx1, ISG15 and IFITM3 in canine tracheal explants infected with CIV(H3N2), CIV(H3N8), IAV(H1N1), AIV(H3N8), EIV(H3N8), SIV(H1N1), AIV(H5N1), BIV(H5N1) and mock-infected controls. Positive immunostaining is shown in brown. Scale bar represents 50 μm. B. Bar plots showing quantification of staining signal by Mx1, ISG15 and IFITM3.

To determine if IFN-mediated responses would render epithelial cells refractory to IAV infection, effectively acting as a barrier against secondary viral infection and hindering reassortment, we treated canine tracheal explants with canine IFN-⍰(_c_IFN-⍰and infected them 24 hours later (when tissues mounted a strong innate immune response, Supplementary Figure 4) with CIV(H3N2) and IAV(H1N1). We chose these viruses because they exhibited the highest replication rates (Figure 2). Figure 8A shows that pre-treatment with _c_IFN-⍰did not affect the growth kinetics of CIV(H3N2). In contrast, IAV(H1N1) replication was significantly reduced at 24 hpi but not at later timepoints (albeit a lower replication trend was observed, Figure 8A). Immunostaining showed double-positive cells to NP and Mx1 (Figure 8B) for both viruses, indicating that CIV(H3N2) and IAV(H1N1) can infect cells that had mounted an antiviral response. These results show that human and canine IAVs can overcome IFN-mediated responses in the canine respiratory epithelium.

**Figure 8.**
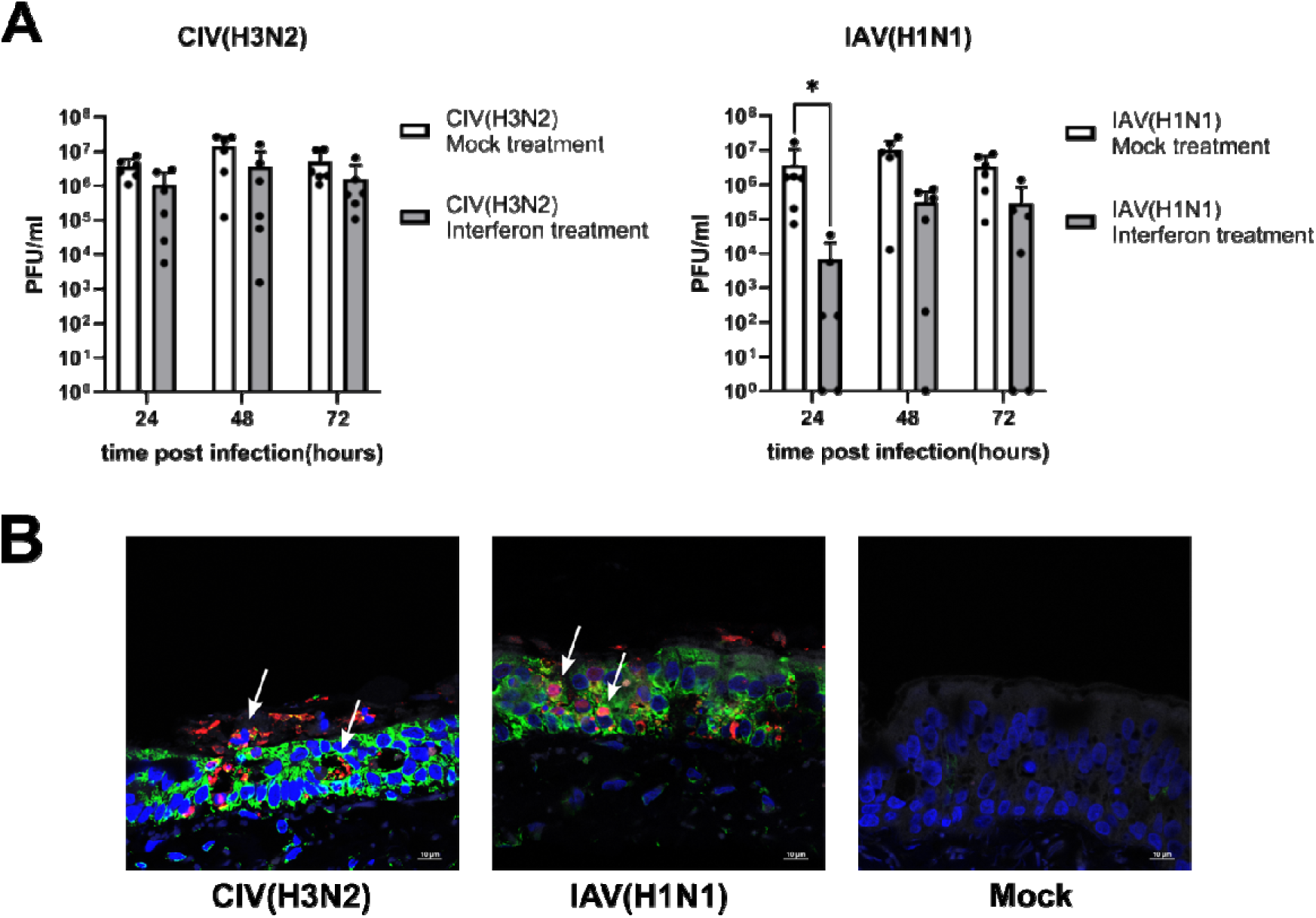
A. Growth kinetics of CIV(H3N2) and IAV(H1N1) in explants derived from canine tracheas that had been pretreated with canine IFN-Ö for 24 hours. Viral titres at individual timepoints are shown in white columns as solid black dots. Each dot represents an individual measurement, and bars represent the mean of six values. Data show combined titres from three independent experiments. B. Representative images of histological sections of explants pre-treated with canine IFN alpha, infected 24 hours later with the indicated viruses, and stained with antibodies against IAV NP (red), and Mx1 (green). Nuclei are stained with Hoechst and shown in fluorescent blue. Arrows indicate infected cells expressing Mx1.

### IAV infections are detected in UK dogs

To determine if domestic dogs are commonly exposed to IAVs we performed and ELISA that detects antibodies against IAV NP on 701 canine residual biochemistry serum samples collected between 2022 and 2023 by the Veterinary Diagnostic Laboratory of the School of Veterinary Medicine of the University of Glasgow (Supplementary Table 3). A total of 9 samples (1.3%) were positive. As CIV does not circulate in the UK nor dogs are vaccinated against CIV, these results suggest that seropositive dogs were exposed to IAV as a result of either external introductions or spillover infections.

## DISCUSSION

Viral emergence is the result of dynamic ecological and evolutionary processes that occur at different scales (from individual cells to multi-host ecosystems) and are inextricably linked. Understanding the relative role of those processes is key to controlling future pandemics. The current panzootic caused by avian-origin H5N1 IAVs and the emergence of a bovine influenza lineage highlight the importance and timeliness of this issue.

Here we focused on the dog as both a potential recipient and/or source of interspecies influenza infections. From an ecological perspective, dogs comprise a large population (∼1 billion^30^) and the high rates of direct contact with humans provide abundant opportunities for bidirectional interspecies transmission. From a virological standpoint, three distinct CIVs emerged in the last 25 years, and each cross-switching event involved a virus from a different species (i.e. horses^12^, birds^13^, and pigs^14^).

To characterise the canine respiratory glycome and the susceptibility and tropism of IAVs we used canine tracheal explants cultured at an air-liquid interface. This experimental system largely represents the natural site of infection in dogs as typical influenza associated pathology includes tracheitis and bronchitis^31^. Additionally, explant cultures are routinely used to study other respiratory viruses ^32,33^.

Virus attachment and entry are mediated by interactions between the viral haemagglutinin and sialylated glycans. Glycomic characterisation showed that sialylated glycans are abundant in the canine trachea, as well as the cultured trachea explants used for our analyses – which were highly representative of the fresh tissue. The sialylated glycans terminated in both α2-6 and α2-3 linked sialic acid, which supports the tissues being permissive to human- and avian-adapted IAVs, as well as equine, bovine, swine and canine strains. Interestingly, the human-adapted IAV(H3N2) virus did not infect the explants. This could be explained by the very low relative abundance of glycans containing multiple LacNac repeats, which human H3 has evolved to preferentially bind to ^34–36^. The majority of sialylated N-glycans detected in our analysis contained just one LacNAc, potentially limiting the ability of IAV(H3N2) to infect and replicate in the canine trachea. It is important to note that we used samples sourced from beagle dogs for our analysis, and that glycosylation in the respiratory tract may differ between dog breeds. Indeed, some Asian breeds express N-glycolylneuraminic acid on their erythrocytes, unlike European breeds^37^.

The probability of infection after cross-species viral exposure is key during spillover transmission^8^ and our results show that with the only exception of IAV(H3N2), all the viruses tested replicated successfully in the dog trachea. It is important to note the high efficiency of infection for each independent and technical replicate, even though we used an inoculum dose that was several orders of magnitude lower than those used in other *ex vivo* studies^25,38^ and lower than normal values of IAV shedding by humans^39,40^, horses^41^, wild birds^42^ and cows^10^.

Segment reassortment is a major force in influenza cross-switching^43^. For reassortment to occur, coinfecting IAVs must exhibit the same cellular tropism. As ciliated cells -the most predominant cell type of the tracheal epithelium- are susceptible to infection by canine, human, equine, bovine and avian viruses, we showed that the dog trachea provides a suitable ecosystem for reassortment. Our findings are supported by a recent study showing that experimental coinfections of dog tracheal explants using canine, human, and avian IAVs generated reassortant viruses capable of infecting human cells^44^.

The timing between primary and secondary viral infections is an important factor in reassortment efficiency, and it has been hypothesised that innate immune responses might preclude secondary infections by heterologous IAVs^45^. In fact, the IFN response triggered by primary viral infection has a protective effect against influenza A viruses^46^. However, our results suggest that antiviral responses are unlikely to block infection by CIV(H3N2) and IAV(H1N1) as they can replicate in explants pre-stimulated with IFN. Thus, CIV(H3N2) and IAV(H1N1) have already overcome an important barrier against segment reassortment.

This work provides serological evidence of IAV exposure in UK dogs. This is consistent with multiple studies that have shown evidence of canine exposure to human^23,24,47–51^ and avian IAVs^52–54^. Given the high level of naïve dogs in countries that are free of canine influenza, the size of the dog population (∼12,5 million dogs in the UK alone^55^), the global co-circulation of human, equine, swine and avian IAVs (including panzootic H5N1), and the susceptibility of the dog respiratory tract to infection by heterologous IAVs, it is feasible that novel CIVs will continue to emerge. The global emergence of IAVs in dogs would not only cause a significant animal welfare issue; it could also increase the risk of zoonotic and anthropozoonotic infections and the subsequent risk of novel IAVs emerging in humans. However, it is important to note that structured contact networks in owned dog populations act as an effective barrier against the spread and maintenance of highly adapted CIVs like H3N8^22^ and H3N2^56^. This *population barrier* makes intervention measures to control CIV emergence more effective, as observed in Canada where multiple clusters of H3N2 CIV infections were contained^57^. However, the emergence and spread of novel CIVs could be more likely and difficult to control in dog populations with high proportion of free-roaming animals exhibiting greater heterogeneity and mixing, as illustrated by rabies in Africa^21,58^.

Our results suggest that dogs could act as a bridging species for the emergence of novel IAVs, including the H5N1 2.3.4.4b genotype. Serological studies in North America and Europe show that hunting dogs are routinely exposed to H5N1 IAVs^53,54^. Vaccination of high-risk dog subpopulations could reduce the risk of interspecies infections (avian-to-dog and dog-to-human) and should be considered as preventative measures.

This study has various limitations. Tracheal explants represent a significant portion of the airways, but this experimental system does not include the upper respiratory tract, the bronchial tree, nor the pulmonary parenchyma, which are likely to exhibit different virus/host interactions due to the presence of different cell types. Further, explants lack systemic responses that are normally mounted during viral infections. Despite these limitations, our results are consistent with observations reported in field studies^59,60^ and experimental *in vivo* infections^26,27^.

In sum, influenza spillover and emergence require a succession of processes that occur at different scales^8^. As the canine respiratory tract is a suitable ecosystem for IAV infection and reassortment and dogs routinely interact with multiple species that support endemic IAVs, dogs could facilitate the flow of virus between sympatric species. Targeted approaches to reduce the risk of IAV infections in dogs should be part of preparedness efforts to control influenza cross-species switching.

**Table 1.** Viruses used in this study.

| **Virus name** | **Abbreviated name** | **Subtype** |
| --- | --- | --- |
| A/Norway/3275/2018* | IAV(H3N2) | H3N2 |
| A/Norway/3433/2018<sup>¶</sup> | IAV(H1N1) | H1N1 |
| A/ruddy<br>shelduck/Mongolia/963V/2009* | AIV(H3N2) | H3N8 |
| A/canine/New York/dog23/2009<sup>¶</sup> | CIV(H3N8) | H3N8 |
| A/canine/Illinois/11613/2015<sup>¶</sup> | CIV(H3N2) | H3N2 |
| A/equine/South Africa/4/2003<sup>¶</sup> | EIV(H3N8) | H3N8 |
| A/Swine/Belgium/2/79 | SIV(H1N1) | H1N1 |
| A/dairy cow/Texas/24-008749-001-original/2024<sup>¶</sup> | BIV(H5N1) | H5N1 |
| A/chicken/England/085598/2022<sup>¶</sup> | AIV(H5N1) | H5N1 |

<sup>¶</sup> Viruses rescued by reverse genetics. \*Virus isolates.

## MATERIALS AND METHODS

### Ethics statement

The use of animal tissues and canine serum samples was approved by the Ethics Committee of the School of Veterinary Medicine of the University of Glasgow (ethics approvals EA26/22 and EA51/24, respectively) and conducted in accordance with the 3Rs principles. As no regulated procedures were carried out on animals by the researchers involved in this study, a Home Office license was not required. Experiments using recombinant viruses were approved by the genetic manipulation safety committee at the University of Glasgow (GM223), and the Health and Safety Executive of the United Kingdom.

### Viruses

An isolate of A/Norway/3275/2018 (H3N2) was provided by the World Influenza Centre. A/ruddy shelduck/Mongolia/963V/2009^61^ and A/canine/New York/dog23/2009 (H3N8)^62^ have been previously described. Plasmids to rescue A/canine/Illinois/11613/2015 (H3N2) were kindly provided by Luis Martinez Sobrido^63^, plasmids to rescue A/England/195/2009 were previously described^64^. A/equine/South Africa/4/2003 (H3N8) RNA was extracted from virus stocks and subcloned into pHW2000 plasmids following RT-PCR. Plasmids to rescue A/Norway/3433/2018 (H1N1), A/dairy cow/Texas/24-008749-001-original/2024, A/chicken/England/085598/2022 and A/swine/Belgium/2/79 were chemically synthesised and cloned into pHW2000.

#### Virus rescue

Viruses were generated as follows: ∼2 × 10^6^ HEK-293T cells were transfected with reverse genetics plasmids (one plasmid for each virus genomic segment). Sixteen hours post-transfection, culture medium was changed to infection medium (serum-free DMEM supplemented with 1 μg/mL TPCK-trypsin [Sigma]). No TPCK was added for highly pathogenic IAVs. The supernatant of transfected cells was collected 48 hours later and used to infect MDCK or MDCK-SIAT1^65^ cells that were maintained in infection medium until the presence of extensive cytopathic effect. At this stage the supernatant was collected, clarified, aliquoted and stored.

#### Tissues and explant preparation

Dog tracheas were collected from healthy Beagles humanely euthanised for reasons unrelated to this study, following their use as controls in other research at Charles River Laboratories. Tissue harvest, transport and preparation have been previously described^62^. The only modification to the original protocol was the use of round 5 mm-wide biopsy punches (Covetrus) to cut individual explants. Explants were maintained at 37°C, 5% CO2, and 95% humidity for up to 5 days.

#### Cells

MDCK, MDCK-SIAT1 and HEK-293T cells were cultured in DMEM (Gibco) supplemented with 10% (v/v) foetal calf serum (Gibco), and 1% non-essential amino acids (NEAA; Gibco Life Technologies, UK). at 37°C with 5% CO2.

#### Glycomic analysis of tissues and tissue explants

Both the canine trachea and trachea explants (five biopsy punches) were washed multiple times with ice cold PBS, then processed as previously described^66^. For the canine trachea, the internal epithelial layer was scraped with a scalpel into ice cold PBS. The samples were sonicated to lyse the cells, then samples were dialysed and lyophilised. Reduction and carboxymethylation reactions were carried out on the samples, which were then treated with trypsin (Sigma) and the resultant glycopeptides were purified by Sep-Pak C18 chromatography. PNGase-F (Roche) was used to release the N-Glycans which were subsequently purified by Sep-Pak C18 chromatography. O-glycans were released by a reductive elimination reaction with potassium borohydride (KBH_4_) and potassium hydroxide (KOH) then subsequently purified using a desalting column of Dowex beads as well as Sep-Pak C18 chromatography. Glycans were then permethylated and analysed using a 4800 MALDI-TOF mass spectrometer (Applied Biosystems). The N-glycans were treated with Sialidase-S or Sialidase-A (Agilent Technologies) to determine the sialic acid linkage configurations. These samples were then permethylated prior to analysis. For both the canine trachea and canine trachea explants, 2 biological repeats were carried out.

Data Explorer (Applied Biosystems) was used to visualise MS and MS/MS data and [M+Na]^+^ molecular ions were annotated manually with the assistance of GlycoWorkbench^67^. Annotations were based on glycan composition and knowledge of biosynthetic pathways. MS/MS analysis was used to confirm predicted assignments and Adobe Illustrator was used to prepare the figures. Glycomics data was quantified in R as previously described^68^. This program was used to detect glycans according to their isotopic peak pattern and calculate the relative intensity across the isotope cluster. The proportions of α2-3 and α2-6 linked sialic acid were then calculated using R^69^. First, the relative intensity of each glycan was normalised to the sum of the relative intensities of all the glycans in the sample. For sialylated glycans, normalised intensities were then weighted based on the number of sialic acid residues present on each glycan. The proportion of α2-6 linked sialic acid was determined by summing the weighted normalised intensities of sialylated molecular ion peaks remaining after sialidase-S digestion. The proportion of α2-3 linked sialic acid was calculated by subtracting the weighted normalised intensities of sialylated molecular ions in the sialidase-S sample from the weighted normalised intensities of sialylated molecular ions from the untreated sample (Supplementary Data File 1).

#### Explant infections

Individual explants were infected apically with 200 plaque forming units (PFUs) of each virus in a total volume of 5 μl. Infection medium without virus was deposited on the surface of mock-infected explants. Infected explants were harvested at defined times post infection and vortexed in DMEM for 10 minutes. Supernatant was aliquoted and stored at -80°C until titration. Each infection was performed in three independent experiments and consisted of two technical replicates.

### Explant interferon treatment

#### Canine IFN-⍰recombinant protein

(Kingfisher Biotech, RP0463D) was dissolved in dH O at a concentration of 40 mg/mL to create the stock solution. The stock solution was then diluted in DMEM to a final concentration of 0.4 µg/mL, producing the working solution. 5 µL of the working solution was added to each explant 24 hours after preparation. DMEM alone was used for mock-treated explants. Treated explants were collected for histological analysis or infected with IAV 24 hours post-treatment.

#### Virus titrations

Virus titrations were carried out in MDCK and MDCK-SIAT1 cells using standard plaque assays as previously described^70^. For plaque assays 5 x 10^5^ cells/ml (MDCK or MDCK-SIAT1) were seeded (12-well plates) and serial dilutions of virus samples were prepared in infection medium. Virus dilutions were added to each well and incubated for 1 hour at 37°C. Subsequently, 1 ml of overlay media (2× MEM; [Life technologies 11935046], 1.2% Avicel [FMC BioPolymer], trypsin, treated with N-tosyl-L-phenylalanine chloromethyl ketone (TPCK) 1 μg/mL [Sigma T4376]) was added. Plates were incubated at 37°C for 72 hours and fixed with 8% formaldehyde in phosphate buffered solution (PBS) and stained with 0.1% Coomassie Brilliant Blue.

#### Immunostaining

Infected explants were harvested at defined times post-infection, fixed in 10% (v/v) buffered formalin and embedded in paraffin. Sections were stained with haematoxylin and eosin (H&E) to assess tissue morphology. Tissue sections were deparaffinized and rehydrated using standard procedures. Briefly, blank sections were first heat-treated in a microwave using pH 6.0 citrate-buffered solution. Following pretreatment, sections were blocked with 1% BSA and incubated with anti-NP primary antibody overnight at 4°C. Donkey anti-Sheep IgG (H+L) cross-adsorbed antibody, Alexa Fluor™ 594(A 11016, 1:200, Invitrogen) was used as secondary antibody for 1 hour at room temperature. Monoclonal Anti-β-Tubulin IV antibody (Sigma Aldrich, T7941, 1:200), Anti-Mucin 5AC antibody (Abcam, 45M1, 1:200) and anti-p63 antibody (Abcam, EPR5701, 1:200) were used as primary antibodies for cell markers and were incubated overnight at 4°C. EnVision+ Single Reagents (HRP. Rabbit), (Dako, Agilent, UK, K4003). Rabbit anti-mouse IgG Alexafluor 488 (Sigma Aldrich, SAB4600056, 1:200) were used as secondary antibodies. Following the immunostaining, explants were washed with dH O and stained with Phalloidin-iFluor 647 Reagent (Abcam, ab176759, 1:500) for 30 minutes. For nuclear staining, slides were treated with Hoescht 33342 solution for 30 minutes, then mounted with ProLong™ Gold Antifade Mountant(Invitrogen, P36935). Images were captured using an Olympus BX51 microscope.

For immunohistochemistry, tissue sections were deparaffinized and rehydrated using standard procedures. Antigen retrieval was performed in citrate buffer, followed by heating. To quench endogenous peroxidase activity, sections were incubated in 3% H O for 10 minutes. Sections were then incubated overnight at 4°C with the following primary antibodies, diluted in 1% BSA: Anti-MX1 Mouse Monoclonal antibody (Clone M143; dilution 1:1000; provided by Georg Kochs), rabbit anti- IFITM3, 1:750 (Proteintech, 11714-1-AP) and rabbit anti-ISG15, 1:1000 (Proteintech, 15981-1-AP). Biotin-SP-conjugated AffiniPure Donkey anti-Sheep IgG(713-065-003, 1:100, Jackson immune research), biotinylated secondary antibodies (anti-mouse IgG(Merck, 1:500, AP181B) and anti-rabbit IgG, 1:500 (Stratech Scientific, 711-065-152) were used as secondary antibodies for 1 hour, followed by extravidine peroxidase (1:100; Merck, E2886-1ML) incubation for 1 hour. Liquid DAB (3,3’-Diaminobenzidine) was applied for 3 to 6 minutes, and slides were counterstained with Mayer’s hematoxylin. Finally, slides were dehydrated through an ascending alcohol series and xylene before being cover slipped.

#### Lectin staining

Sections were deparaffinized after paraffin embedding, rehydrated, and retrieved in 0.01M EDTA for 30 minutes at 95°C. After cooling to RT, sections were washed three times with PBS and blocking with 1% BSA in room temperature 30 minutes, and incubated with 20 μg/mL Fluorescein-conjugated MAL-I or CY5-conjugated SNA (Vector Laboratories) overnight at 4°C. After lectin incubation, sections were treated with streptavidin conjugated with FITC (Invitrogen) and incubated for 1 hour at 4°C. Finally, slides were stained with Phalloidin-iFluor 647 Reagent (Abcam, ab176759, 1:500) for 30 minutes. For nuclear staining, slides were treated with Hoescht 33342 solution for 30 minutes, then mounted with ProLong™ Gold Antifade(Invitrogen, P36935). Images were captured using an Olympus BX51 microscope.

## Quantification of infected cells

The number of IAV infected ciliated and goblets cells were counted. Briefly, to determine the number of infected ciliated cells, the total number of NP-positive cells as well as double positive cells for IAV NP/β-tubulin were manually counted. For goblet cells a similar approach was taken but counting the total number of Mucin 5AC single-positive cells as well as double positive (IAV NP/Mucin 5AC) cells. Two sections per donor were counted.

## Digital Pathology/Image Analysis

Mx1, ISG15 and IFITM3 IHC staining was quantified using HALO v4.1 (Indica Labs). Briefly, digitised whole slide scans were annotated to indicate the region of interest and exclude artefacts, then segmented by a HALO AI DenseNet V2 mask classifier trained to identify respiratory epithelium from surrounding submucosa, mucus, cell debris and slide glass. ISG quantification was performed on areas positively identified as epithelium using the HALO Area Quantification v3.0.1 algorithm, which designates DAB-stained pixels as negative or weak, moderate, or strong positive based on minimum optical densities (weak positive: 0.15; moderate positive: 0.2165; strong positive: 0.3835). All epithelium identification and IHC staining quantification was manually checked following analysis.

### Serological testing

Enzyme linked immunosorbent assay (ELISA) was performed using IDEXX Influenza A test Kit to detect antibodies to influenza A.

The assay was performed following the manufacturer’s instructions, using positive and negative controls provided with the kit to validate each assay run. Samples were diluted 1:10 in dilution buffer. Briefly,100 µl of diluted samples and undiluted controls were added to corresponding wells of the ELISA plates and incubated at room temperature for an hour. Following incubation, plates were washed 3 times with freshly prepared wash buffer (diluted 1:10). An anti-influenza A nucleoprotein monoclonal antibody enzyme conjugate was added to each well of the ELISA plate. The plate was then incubated at room temperature for 30 minutes, washed to remove excess unbound conjugate, followed by the addition of 100ul/well 3,3′, 5,5 ′ -tetramethylbenzidine (TMB) substrate. After 15 minutes of incubation at room temperature the colour development was terminated by adding 100 µl of stop solution. The optical density was measured at 650 nm in Multiskan FC plate reader (Thermo Fisher). The assay was considered valid if the mean of the negative control was greater than 0.60 and the mean of the positive control divided by the value of the negative control mean was less than 0.50. For each sample the index value was determined by calculating sample absorbance divided by absorbance of negative control mean. Samples with index values of 0.6 or less were considered as positive.

### Statistical analysis

Differences between viral titres were analysed using Mann-Whitney U tests in Graph Pad 10. P values <0.05 were considered significant.

## ACKNOWLEDGMENTS

We thank Dr Christoph Blau (Charles River) for providing tissues. We also thank Rebecca Ross for technical assistance. We thank Lynn Stevenson and Frazer Bell for support with histological preparations. We gratefully acknowledge all data contributors (i.e. the authors and their originating laboratories responsible for obtaining the specimens, and their submitting laboratories for generating the genetic sequence and metadata and sharing via the GISAID Initiative^71^, on which this research is based. All submitters of the data may be contacted directly via the GISAID website (https://www.gisaid.org).

## SUPPLEMENTARY TABLES

**Supplementary Table 1.** Quantification of IAV-infected ciliated cells.

| NP positive ciliated cell/ NP positive cell | CIV H3N2 | CIV H3N8 | IAV H1N1 | AIV H3N8 | SIV H1N1 | EIV H3N8 | AIV H5N1 | BIV H5N1 |
| --- | --- | --- | --- | --- | --- | --- | --- | --- |
| Donor 1 | 10/24 | 44/59 | 4/14 | 2/8 | 1/3 | 1/3 | 1/5 | 2/6 |
|  | 7/22 | 41/62 | 4/17 | 3/12 | 1/4 | 0/4 | 1/3 | 2/5 |
| Donor 2 | 47/78 | 43/85 | 28/48 | 31/46 | 0/3 | 6/14 | 5/16 | 3/17 |
|  | 68/98 | 42/70 | 24/53 | 25/54 | 0/1 | 5/12 | 3/20 | 2/13 |
| Donor 3 | 6/28 | 36/66 | 69/86 | 4/13 | 0/2 | 1/4 | 1/15 | 0/10 |
|  | 8/26 | 28/63 | 70/83 | 2/10 | 3/7 | 1/5 | 2/17 | 0/8 |

**Supplementary Table 2.**
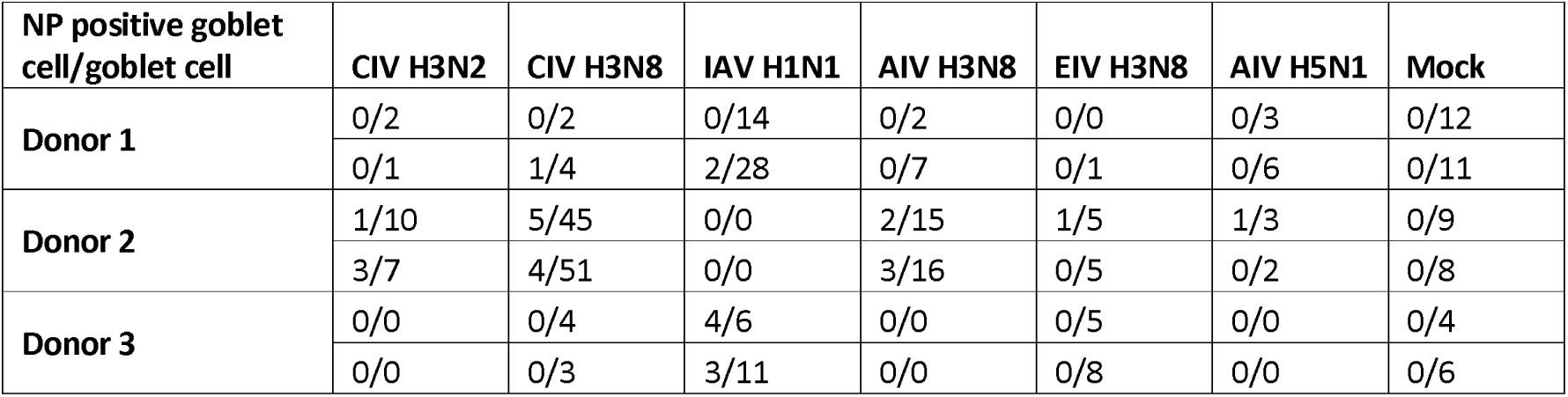
Quantification of IAV-infected goblet cells.

## SUPPLEMENTARY FIGURES

**Supplementary Figure 1.**
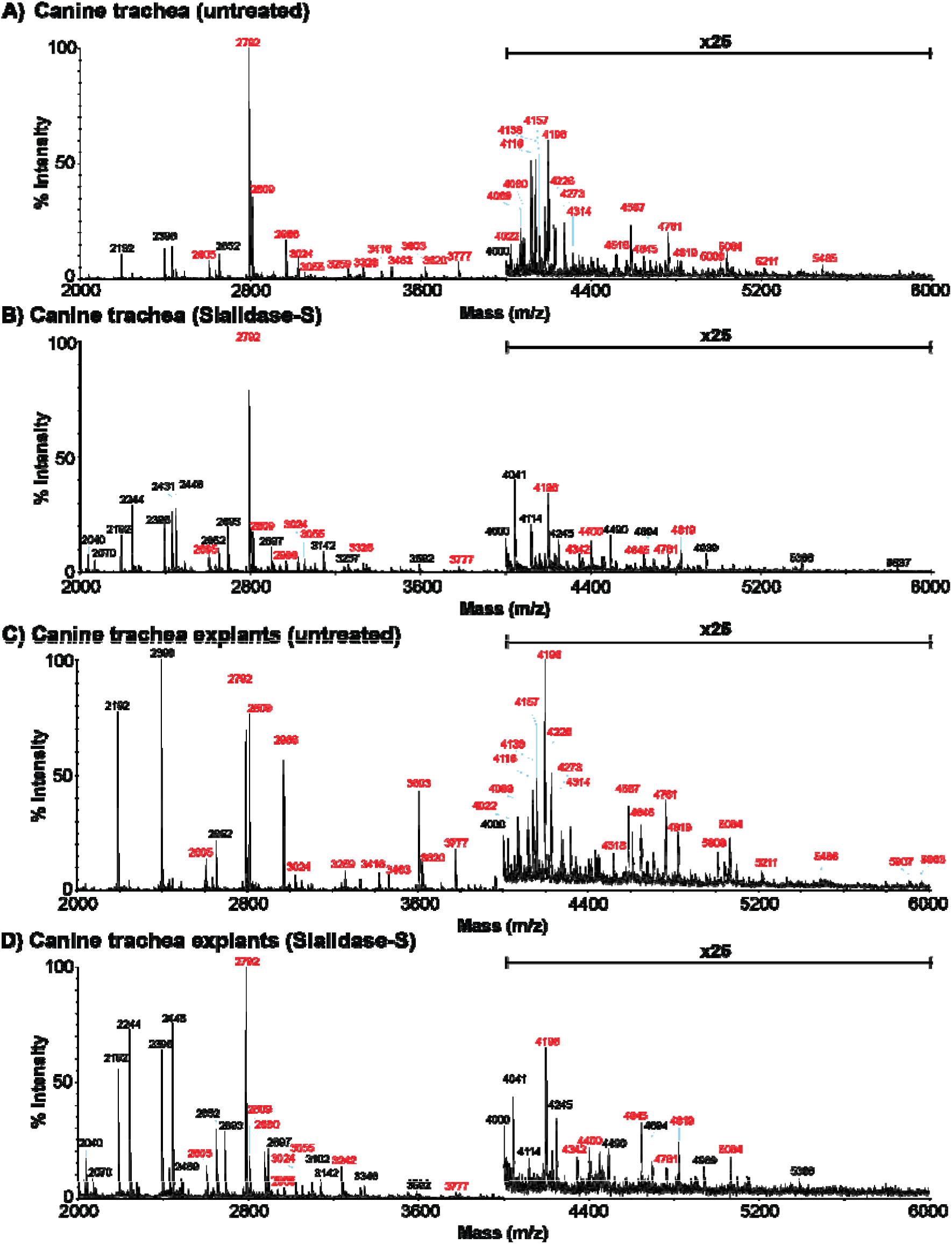
MALDI-TOF spectra of permethylated N-glycans from canine tracheas and trachea explants following sialidase digestions. Annotated MALDI-TOF spectra of A) N-glycans from canine trachea, B) N-glycans from canine trachea treated with sialidase-S, C) N-glycans from canine trachea explants and D) N-glycans from canine trachea explants treated with sialidase-S. Annotations show [M + Na]^+^ molecular ions and include major structures which were unsialylated (black) and sialylated (red).

**Supplementary Figure 2.**
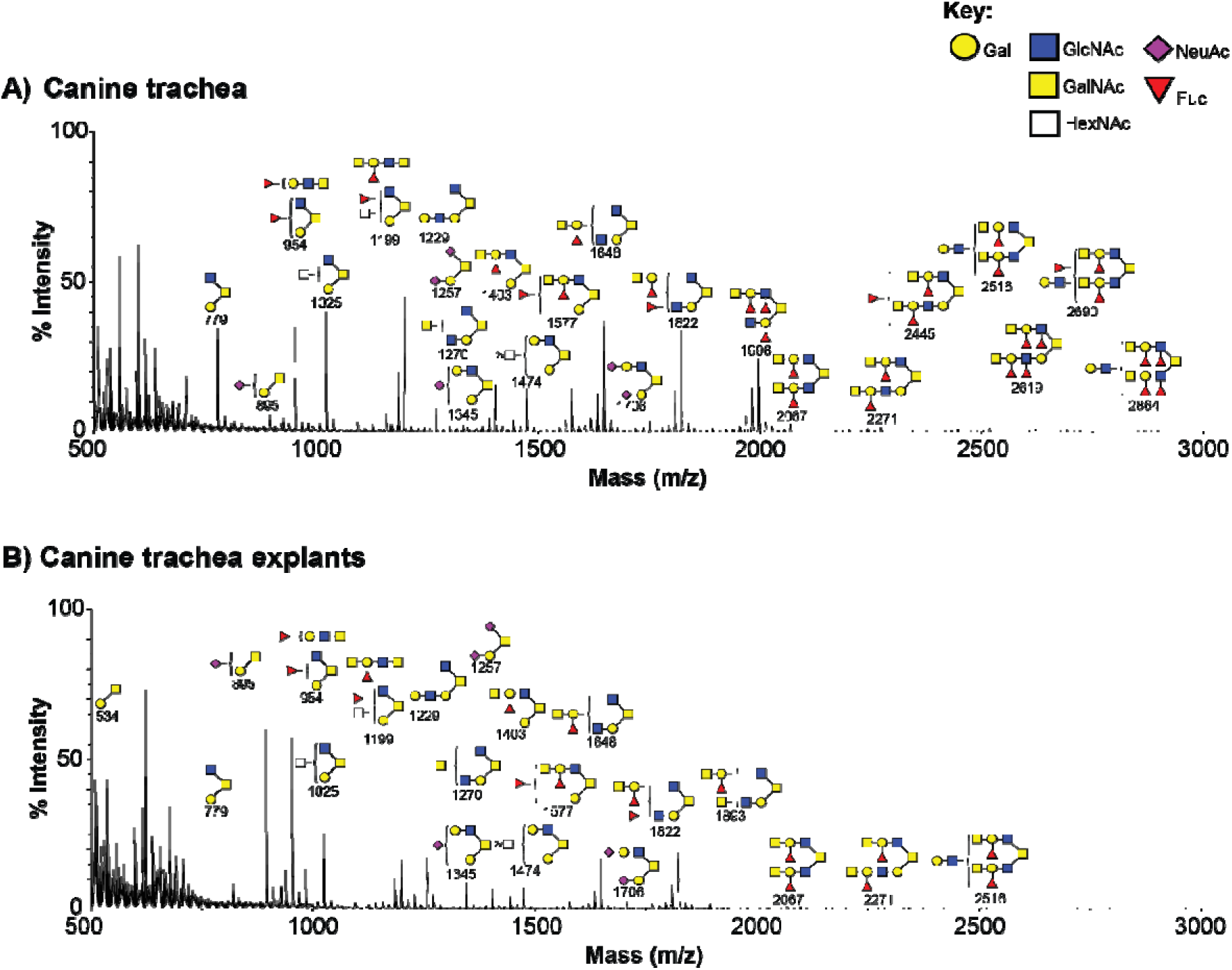
MALDI-TOF spectra of permethylated O-glycans from canine tracheas and trachea explants. Annotated MALDI-TOF spectra of O-glycans from A) canine trachea and B) canine trachea explants. Annotations show [M + Na]^+^ molecular ions. Peak annotation is based on composition, biosynthetic knowledge and MS/MS analysis.

**Supplementary Figure 3:**
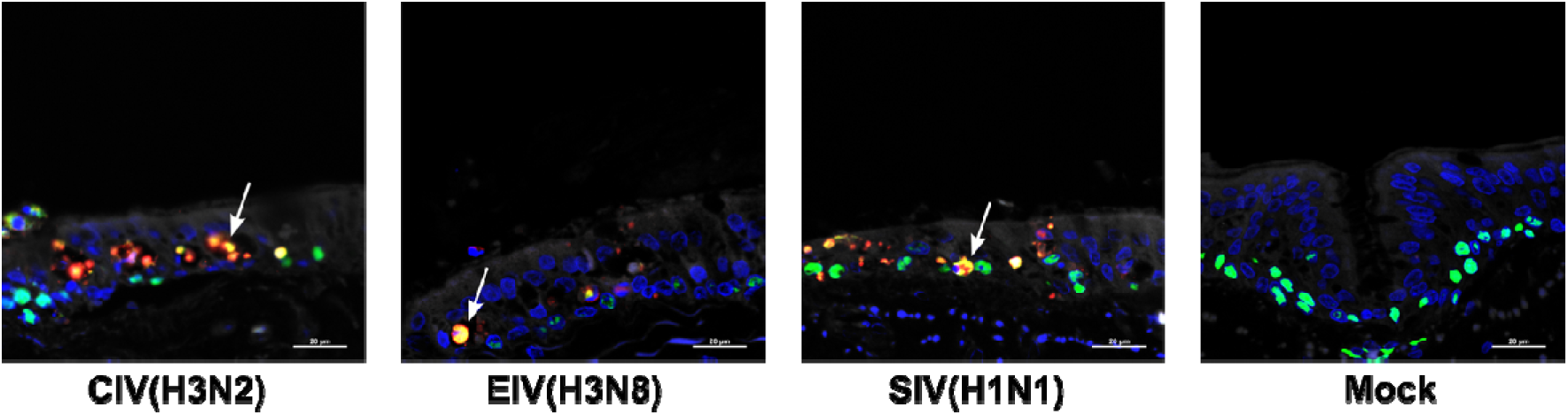
Cellular tropism of IAVs. Representative images of immunostained sections derived from canine tracheal explants infected with CIV(H3N2), EIV(H3N8), and SIV(H1N1) and mock-infected controls. Viral NP is shown in fluorescent red and indicates infected cells, whereas fluorescent green shows immunostaining of p63, indicating basal cells. Arrows indicate infected basal cells.

**Supplementary Figure 4:**
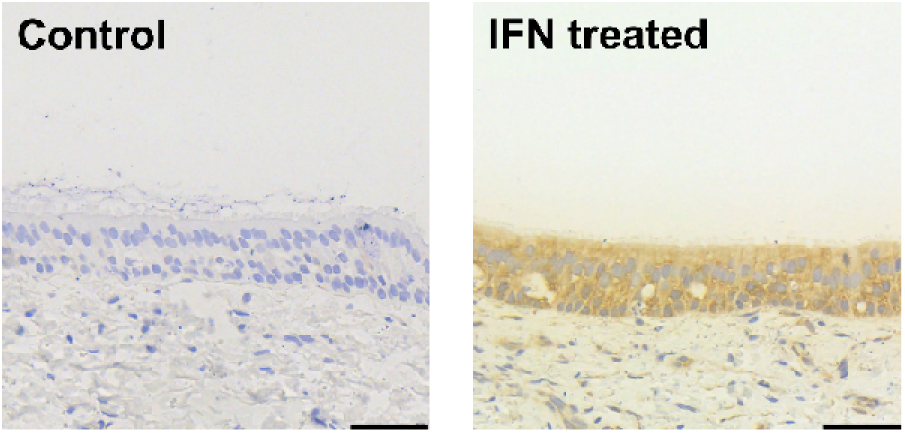
Expression of Mx1 in canine tracheal explants treated with canine interferon for 24 hours. Positive immunostaining is shown in brown. Scale bar represents 50 μm.

**Supplementary Figure 5:**
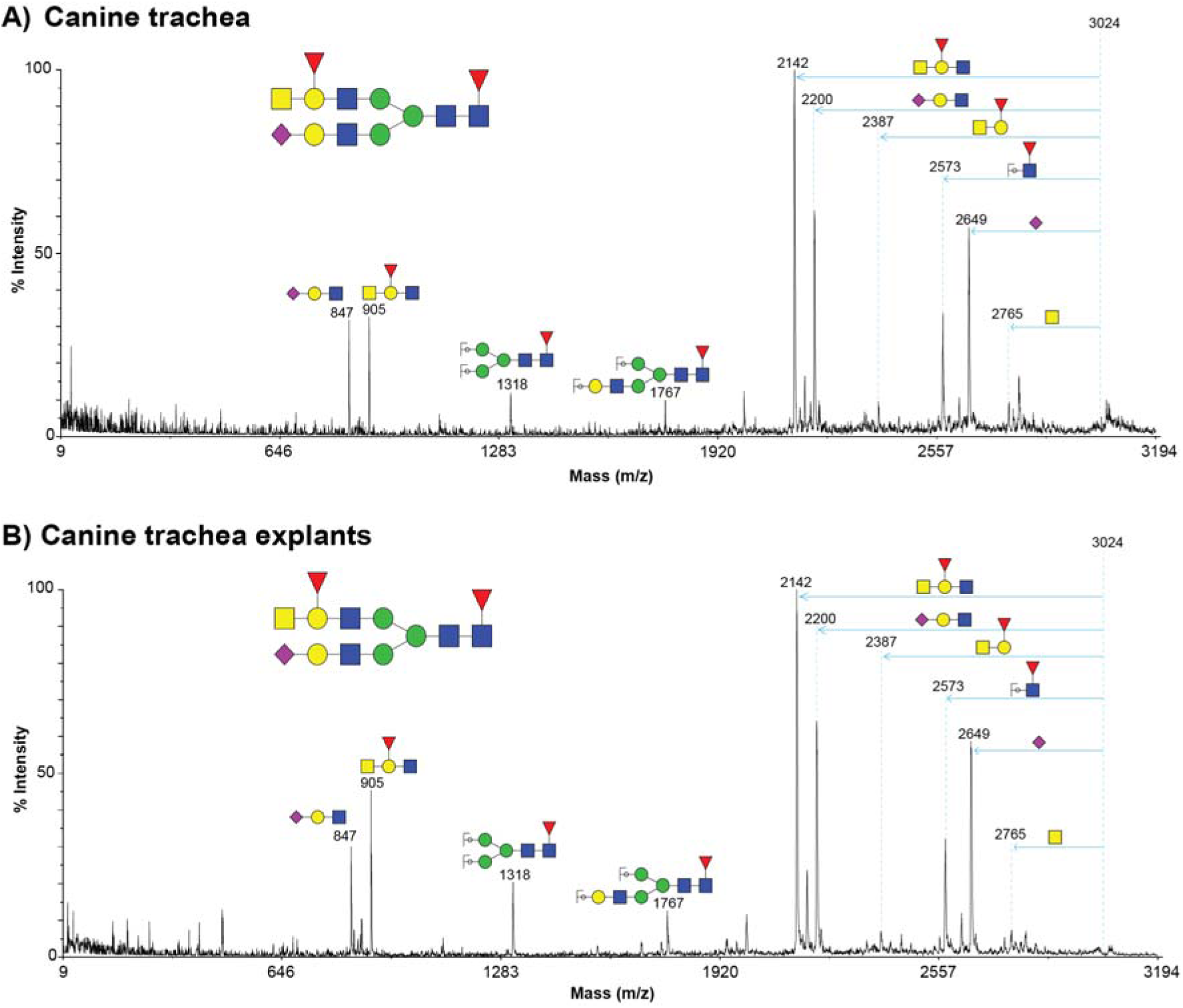
MALDI TOF/TOF MS/MS spectra of m/z 3024. MALDI TOF/TOF MS/MS spectrum of *m/z* 3024 from A) canine trachea and B) canine trachea explants showing example MSMS data of a sialylated N-glycan with an ABO blood group A epitope. Annotations show [M + Na]^+^ fragment ions. Horizontal arrows correspond to the loss of the indicated fragment from the molecular ion.

